# Distinct features of the regenerating heart uncovered through comparative single-cell profiling

**DOI:** 10.1101/2023.07.04.547574

**Authors:** Clayton M. Carey, Hailey L. Hollins, Alexis V. Schmid, James A. Gagnon

## Abstract

Adult humans respond to heart injury by forming a permanent scar, yet other vertebrates are capable of robust and complete cardiac regeneration. Despite progress towards characterizing the mechanisms of cardiac regeneration in fish and amphibians, the large evolutionary gulf between mammals and regenerating vertebrates complicates deciphering which cellular and molecular features truly enable regeneration. To better define these features, we compared cardiac injury responses in zebrafish and medaka, two fish species that share similar heart anatomy and common teleost ancestry but differ in regenerative capability. We used single-cell transcriptional profiling to create a time-resolved comparative cell atlas of injury responses in all major cardiac cell types across both species. With this approach, we identified several key features that distinguish cardiac injury response in the non-regenerating medaka heart. By comparing immune responses to injury, we found altered cell recruitment and a distinct pro-inflammatory gene program in medaka leukocytes, and an absence of the injury-induced interferon response seen in zebrafish. In addition, we found a lack of pro-regenerative signals, including nrg1 and retinoic acid, from medaka endothelial and epicardial cells. Finally, we identified alterations in the myocardial structure in medaka, where they lack embryonic-like primordial layer cardiomyocytes, and fail to employ a cardioprotective gene program shared by regenerating vertebrates. Our findings reveal notable variation in injury response across nearly all major cardiac cell types in zebrafish and medaka, demonstrating how evolutionary divergence influences the hidden cellular features underpinning regenerative potential in these seemingly similar vertebrates.

## INTRODUCTION

Myocardial infarction (MI), commonly known as a heart attack, contributes significantly to human morbidity and mortality^1^. During an MI, a blockage in a coronary artery cuts off blood flow to the heart muscle causing cell death and the eventual formation of a non-contractile scar. In adult mammals, including humans, this scar is permanent and impairs cardiac function^2^. In contrast, many types of fish and amphibians possess the remarkable ability to clear cardiac scar tissue and regrow damaged muscle as adults^3, 4^. These observations have sparked intensive studies of regenerating species in hopes of discovering evolutionarily conserved mechanisms to enable regeneration in humans^5^. Such comparative studies are confounded, however, by the large evolutionary divergence between mammals and regenerating vertebrates. This distant evolutionary relationship results in often unclear gene orthology to mammals and manifests in the distinct simplified heart anatomy of fish and amphibians. Thus, despite many advances, the precise molecular, cellular, and genetic factors that enable some animals to regenerate as adults remain incompletely defined.

Zebrafish have emerged as a powerful model for studying adult heart regeneration^6, 7^. Experimentally induced ventricular cryoinjury is frequently used in zebrafish to mimic infarction events seen in humans^8^. Following injury with a liquid nitrogen-cooled probe, a lesion of necrotic tissue forms, triggering an acute inflammatory response that recruits various immune cell types to the wound^9^. The activities of these immune cells play a crucial role in the subsequent remodeling and regeneration processes in zebrafish. Macrophages and regulatory T cells, in particular, are indispensable for successful regeneration^10, 11^. Additionally, fibroblast cells derived from both the endocardium and epicardium become activated and deposit the collagenous matrix that makes up the scar and stabilizes the injured ventricle^12, 13^. In zebrafish, activated fibroblasts also provide critical signals that foster a regenerative niche by promoting neovascularization of the wound area and dedifferentiation and proliferation of existing cardiomyocytes in the wound border zone. This signaling is mediated in part by molecules such as nrg1, secreted from epicardial-derived cells^14^, and retinoic acid, chiefly produced by the endocardial compartment^15^. The synergistic effects of these signals promote the regrowth of coronary vessels and replacement of scar tissue with healthy myocardium. Although these cellular behaviors are well-established in zebrafish, it remains unclear whether non-regenerating species might share some or all of these characteristics. Therefore, comparative studies are still needed to determine which behaviors of immune cells and components of the signaling environment are truly unique to the regenerating heart.

Recent surveys of cardiac regeneration capabilities among different teleost fish species have yielded surprisingly contrasting results, demonstrating that zebrafish cardiac injury responses are not representative of all teleosts. While ventricular regeneration is found in some fish species^16, 17^, several others, including the grass carp^18^, Mexican cavefish^19^, and Japanese medaka^20^, exhibit permanent scarring similar to adult mammals. The cellular and molecular behaviors that distinguish these non-regenerating species from zebrafish, however, have only begun to be characterized at the cellular level. The existence of non-regenerating teleosts offers a unique opportunity to compare and contrast the differing regeneration phenotypes across relatively short evolutionary distances to determine which cellular features are unique to regenerating species. Given that heart regeneration was likely an ancestral trait of teleosts^4, 15^, understanding the evolutionary path that led to the loss of this ability in some species may offer parallel insights into why mammals lose the ability to regenerate as adults.

In this study, we used comparative single-cell transcriptomics to create detailed time-course maps of the cardiac injury response in zebrafish and the non-regenerating Japanese medaka. These fish share similar body and heart anatomy and shared a common teleost ancestor ∼140 million years ago^21^. Our approach revealed key differences in both pre- and post-injury hearts that may be responsible for the contrasting regeneration outcomes. We found differences in immune cell recruitment and behavior, epicardial and endothelial cell signaling, and alterations in the structure and makeup of the myocardium. Overall, our findings shed new light on the factors that coordinate heart regeneration and generate new hypotheses for the mechanisms that underlie the loss of this ability in certain species.

## RESULTS

### Time-resolved atlas of cardiac injury response in zebrafish and medaka ventricles

There have been few direct cross-species comparative studies of cardiac injury thus far^19, 22^. Single-cell RNA sequencing (scRNA-seq) is a powerful approach that enables simultaneous comparison of cellular composition and gene expression across all major cell types in the heart. Therefore, we sought to create a comparative cell atlas of cardiac injury response in zebrafish (*Danio rerio*) and medaka (*Oryzias latipes*) ventricles. We used cardiac cryoinjury to induce comparable injury in the ventricles of both zebrafish and medaka (see Methods). We collected and analyzed uninjured hearts, as well as hearts at 3 and 14 days post-injury (d.p.i.), from both species with several biological replicates at each time point (Figure 1A). Dissected ventricles were dissociated into a single-cell suspension and prepared for scRNA-seq. Consistent with prior studies^8^, cryoinjury-induced lesions in the myocardium and formation of a fibrous clot by 3 d.p.i., with deposition of a collagen-rich scar evident by 14 d.p.i. (Figure 1B).

**Figure 1:**
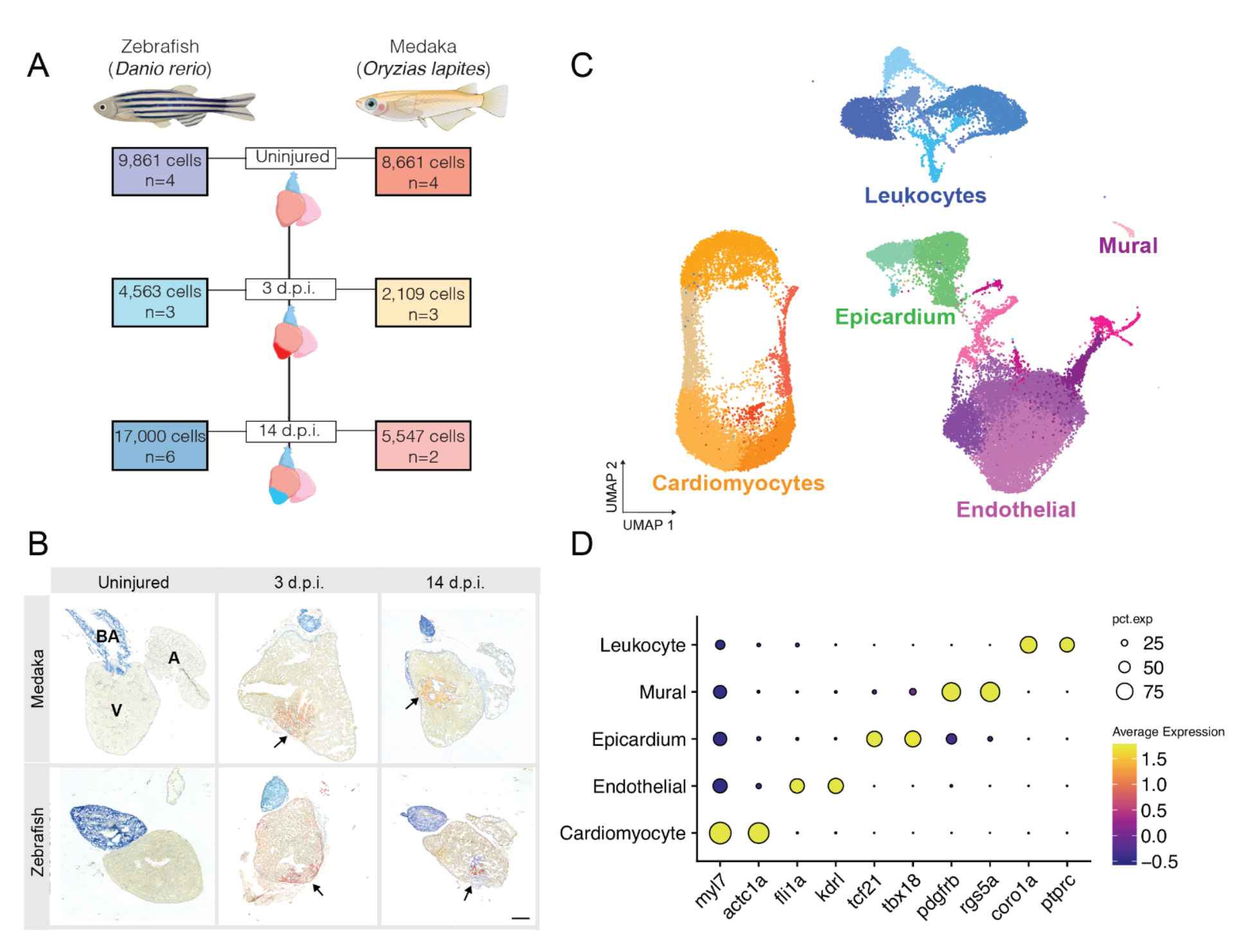
A single-cell atlas of cardiac injury response in zebrafish and medaka. (A) Experimental overview for collection of ventricles and single-cell sequencing. The number of independent samples and total number of quality filtered cells for each time point are indicated. (B) Representative images of Acid-fuchsin orange staining of collagen (blue), fibrin (red), and muscle fibers (tan) in ventricle sections showing cryoinjury-induced fibrin and collagen deposition in both species (arrowheads). Anatomical labels indicate ventricle (labeled V), atrium (labeled A), and bulbus arteriosus (labeled BA). (C) UMAP embedding of all sampled cells from each species and time point integrated into a single dataset. A total of 22 clusters were identified and colored by major cardiac cell type (cardiomyocyte = orange shades, Endothelial/mural = purple shades, epicardial = green shades, leukocyte = blue shades). (D) Gene expression dot plot showing average gene expression of marker genes for cells classified as the indicated cell type. Two marker genes are displayed for each cell type. Dot sizes represent percent of cells expressing the indicated gene (pct.exp), color indicates average scaled gene expression across all cells in the indicated tissue.

A total of 47,741 non-erythroid cells that passed quality control and filtering parameters were obtained across all samples. To improve downstream data integration, gene names were standardized to unify names where one-to-one orthology was supported (Figure S1A). The resulting cells were log normalized and integrated using Seurat v4^23^. Cell integration anchors were calculated using canonical correlation analysis, a technique that benchmarks well for joint-embedding of shared cell types while maintaining species-specific cell types^24^. After integration, dimensionality reduction and clustering yielded 22 cell clusters (Figure 1C). Analysis of marker gene expression revealed five main cell types including cardiomyocytes (marked by *myl7* and *actc1a* expression), endothelial cells (*fli1a*/*kdrl*^25, 26^), epicardial cells (*tcf21*/*tbx20*^27^), vascular mural cells (*pdgfrb*/*rgs5a*^28^), and leukocytes (*coro1a*/*ptprc*^29^) (Figure 1D). Cross-species data integration was effective as both zebrafish and medaka cells were represented in each major cluster (Figure S1B-C).

To facilitate dissemination of the data to the research community, we created a freely available web-based application for exploration of the single-cell dataset^30^. This tool allows users to examine and compare gene expression and cell type compositions across both species at all time points. Access and instructions for usage can be found at https://github.com/clay-carey/medaka_zebrafish_regeneration.

### Medaka lack an endogenous interferon response to injury

We first compared injury-induced immune responses to determine how medaka might respond differently than zebrafish to immunostimulatory signals released after cryoinjury. Interferon signaling can modulate gene expression in various cell types and may promote regeneration in tissues such as the intestine and skeletal muscle^31, 32^. Previous studies have shown that injection of poly(I:C), a synthetic double-stranded RNA species and a potent activator of interferon signaling via TLR interaction, can enhance revascularization and cardiomyocyte proliferation in injured medaka hearts^22, 33^. To explore whether endogenous interferon signaling may be activated in zebrafish after cryoinjury, we examined the expression of 12 genes previously identified as zebrafish interferon-stimulated genes (ISGs) across different tissue types in our dataset^34^. We observed a surge in ISG expression at 3 d.p.i. in the zebrafish endothelial, epicardial, and leukocyte compartments, but not in the myocardium. In contrast, we did not detect ISG upregulation at 3 d.p.i. or 14 d.p.i. in medaka (Figure 3A). Using RNA in situ hybridization of injured zebrafish hearts, we observed expression of *isg15*, a highly conserved ISG across vertebrates, in the wound area co-mingled with *kdrl*^+^ endothelial cells at 3 d.p.i., but not in 14 d.p.i hearts. Consistent with the single-cell data, we detected little *isg15* expression in injured or uninjured medaka hearts (Figure 2B). Examination of *isg15* expression by cell cluster revealed upregulation in most zebrafish endothelial, epicardial, and mural cell clusters at 3 d.p.i., consistent with a systemic interferon response to injury in zebrafish (Figure S2A).

**Figure 2:**
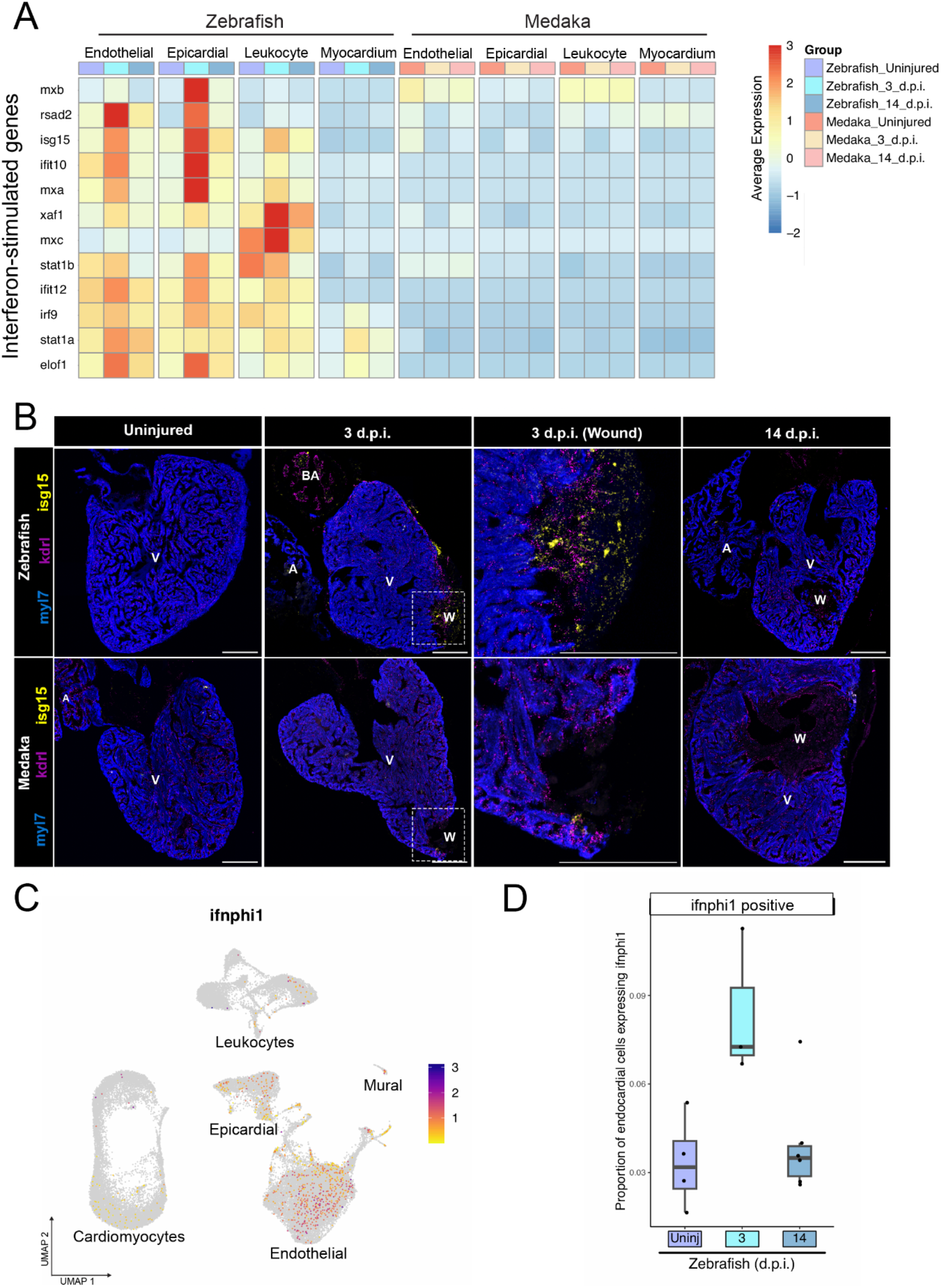
Medaka lack an endogenous injury-induced interferon response. (A) Gene expression heatmap showing scaled average gene expression for 12 interferon-stimulated genes in the indicated species and tissue type at each time point. (B) RNA in situ hybridization of *isg15* (interferon-responding cells), *kdrl* (endothelial cells), and *myl7* (cardiomyocytes) in ventricle cryosections in the indicated species and time point. Scale bar = 200 µm. Anatomical labels: V = intact ventricle, BA = bulbus arteriosus, W = wound area. (C) Gene expression feature plot for *ifnphi1* across all zebrafish cardiac cell types, color scale = expression level. (D) Quantification of proportion of zebrafish endothelial cells expressing *ifnphi1* at each time point.

**Figure 3:**
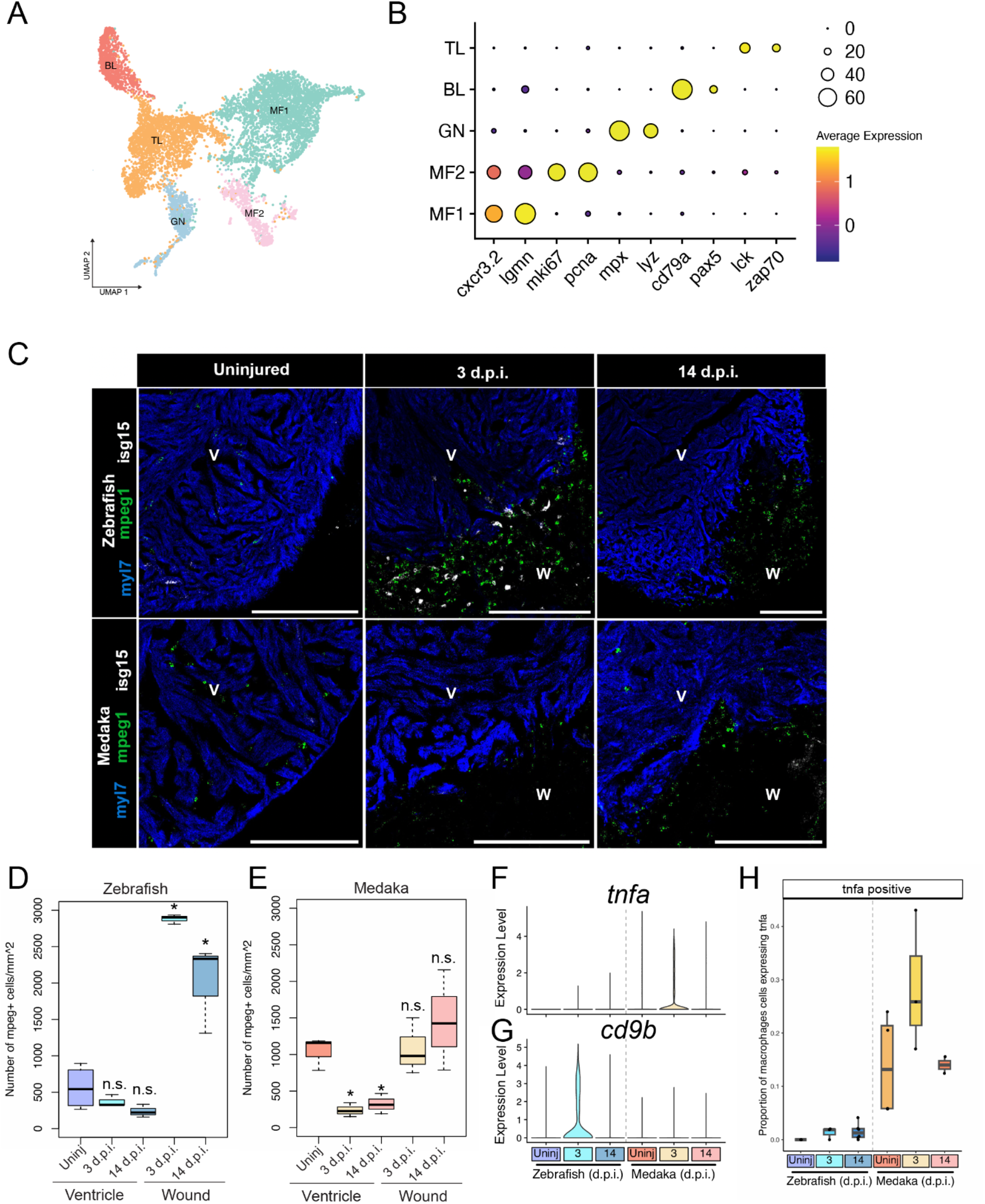
Medaka display altered tissue-resident and injury-responsive immune cell populations. (A) UMAP embedding and sub-clustering of all leukocytes, with cells classified as T lymphocytes (TL), B lymphocytes (BL), Granulocytes (GN), or macrophages (MF). (B) Gene expression dot plot of marker genes for each immune cell cluster. (C) RNA in situ hybridization of *mpeg1.1* (macrophages), *isg15* (interferon response), and *myl7* (cardiomyocytes) in ventricle cryosections in the indicated species and time point. Scale bar = 200 µm, anatomical labels: V = intact ventricle, W = wound area. (D-E) Quantification of number of macrophages per mm^2^ in either the intact myocardium (ventricle) or wound area (wound) in zebrafish (D) or medaka (E) * indicates a p value < 0.05 using a t-test comparing with uninjured ventricle. (F-G) Gene expression violin plots from all macrophages in the indicated time point and species for *tnfa* (F) and *cd9b* (G). (H) Quantification of the proportion of macrophages expressing *tnfa* at each time point in each species.

To investigate the source of post-injury interferon signals in zebrafish, we examined the expression of each interferon isoform annotated in the zebrafish genome. Our analysis revealed that only the *ifnphi1* isoform was expressed in a significant number of cells, and it was primarily restricted to endothelial cells (Figure 2C, Figure S2B). After cell quantification, we found that there was an approximately threefold increase in the number of *ifnphi1*^+^ endothelial cells following injury, which returned to baseline levels by 14 d.p.i. (Figure 2D). These results indicate that endothelial cells are a major source of interferon signaling after injury in zebrafish, possibly in response to damage-associated molecules, which may include immunostimulatory nucleic acids released from necrotic cells^35^. In contrast, medaka hearts do not seem to respond to these damage-associated signals with endogenous interferon signaling, highlighting a critical difference in injury response of these two teleost species.

### Medaka display altered tissue-resident and injury-responsive immune cell populations

Immune cells play a critical role in facilitating immediate injury responses and in modulating tissue regeneration, and may represent a key point of phenotypic variation across species^36, 37^. In prior studies, analysis of bulk transcription patterns highlighted the differential expression of immune-related genes in zebrafish and medaka hearts following injury^22^. We further investigated the immune response in these species by examining immune cell populations through the subsetting and re-clustering of all cell clusters identified as leukocytes (Figure 3A). By using marker gene expression, we identified five distinct cell clusters, corresponding to macrophages (*cxcr3.2/lgmn*), proliferating macrophages (*mki67*/*pcna*), T lymphocytes (*lck/zap70*), B lymphocytes (*cd79a/pax5*), and granulocytes (*mpx/lyz*) (Figure 3B).

Macrophages are essential for regeneration in zebrafish, but medaka display altered macrophage recruitment^22^. To further investigate how medaka macrophages respond to injury, we quantified macrophage dynamics in the intact and injured areas of the zebrafish and medaka ventricle with RNA in situ hybridization and imaging of cells expressing macrophage marker *mpeg1.1*. By imaging healthy and injured ventricles, we found that macrophages are highly enriched in the wound area in zebrafish compared to medaka, localizing near cells expressing *isg15* (Figure 3C). Quantification of *mpeg1.1*^+^ cells in zebrafish demonstrated that macrophage density in the wound is approximately 5-fold higher than in uninjured myocardium at 3 and 14 d.p.i., but macrophage density in the unaffected tissue remains constant (Figure 3D). In contrast, the wound area in medaka ventricles had a similar density of *mpeg1.1^+^* cells to the intact uninjured ventricle, but the macrophages appeared to be depleted from intact tissue after injury in medaka (Figure 3E). These results show that medaka have few signs of post-injury macrophage recruitment or proliferation, while zebrafish macrophage populations are highly expanded in the wound area.

We next compared gene expression in zebrafish and medaka macrophages to determine whether they may have different behaviors in response to injury. Strikingly, medaka macrophages highly upregulated *tnfa* expression at 3 d.p.i. compared to zebrafish, a marker of inflammatory M1-like macrophages (Figure 3F). Conversely, zebrafish macrophages express higher levels of *cd9b* after injury, a marker of antiinflammatory M2-like macrophages^38^ (Figure 3G). Cell quantification from single-cell data showed a slight increase in *tnfa^+^* macrophages in 3 and 14 d.p.i. zebrafish, but uninjured medaka ventricles display a substantially higher population, further increasing after injury (Figure 3H). Our findings show that although medaka have lower macrophage recruitment, they exhibit a more pro-inflammatory response to injury, highlighting a crucial difference in the immune response to injury between the two species.

### Zebrafish and medaka share a partially overlapping fibrotic response to injury

Formation of a fibrotic scar following cardiac injury is an evolutionarily conserved trait among vertebrates^3^. In zebrafish, the scar is chiefly deposited by activated fibroblasts derived from pre-existing epicardial and endothelial cells, but it remains unclear how this process might differ in species that exhibit permanent scarring^13, 39^. We investigated whether medaka might differ in their fibrotic response to injury compared to zebrafish. To identify individual cell types contributing to fibrosis, we first conducted a re-clustering of all cells expressing the endothelial markers *kdrl* and *fli1a*, along with perivascular cells expressing mural cell markers *rgs5a* and *pdgfrb*. This process led to the identification of 11 cell clusters, categorized using marker gene expression as endocardial endothelium (eEC), fibroblast-like endothelial cells (fEC), coronary endothelium (cEC), and lymphatic endothelium (lEC) (Figure 4A-B). Cells in the fEC cluster expressed high levels of collagen isoforms *col1a2* and *col5a1*, as well as *postna* and *twist 1b*, markers of a transition to a mesenchymal activated fibroblast state, but maintained expression of endothelial cell markers *fli1a/kdrl* but not epicardial markers *tcf21/tbx18* (Figure S4A / Figure S4C).

**Figure 4:**
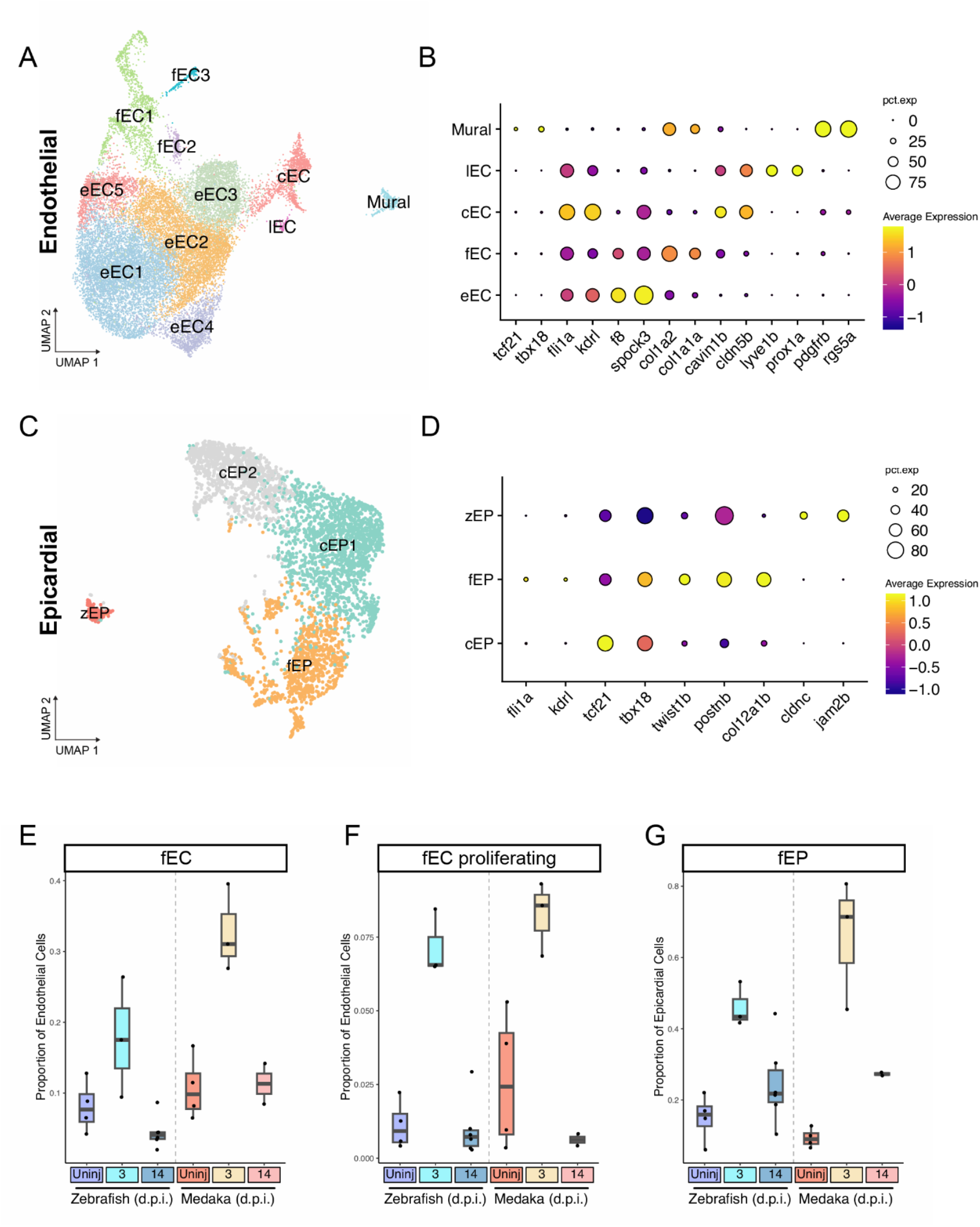
Zebrafish and medaka share a partially overlapping fibrotic response to injury. (A) UMAP embedding of re-clustered endothelial and mural cells classified as either endocardial endothelium (eEC), coronary endothelium (cEC), lymphatic endothelium (lEC), fibroblast-like endothelial cells (fEC), and mural cells (mural). (B) Gene expression dot plot of marker genes for each endothelial cell classification. (C) UMAP embedding of re-clustered epicardial cells classified as canonical epicardial cells (cEP), fibroblast-like epicardial cells (fEP), or zebrafish-specific epicardial cells (zEP). (D) Gene expression dot plot of marker genes for each epicardial cell classification. (E-F) Quantification of proportion of endothelial (E-F) or epicardial (G) cells classified as the indicated cell type.

To investigate the contributions of epicardial-derived cells to the fibrotic response, we re-clustered all cells expressing epicardial-specific markers *tcf21* and *tbx18*, and re-clustered them into four distinct epicardial cell clusters consisting of three cell types: canonical epicardial cells (cECs), fibroblast-like epicardial cells (fEP), and a zebrafish-specific cluster of epicardial cells (zEP) (Figure 4C). The fEP cells are characterized by the expression of epithelial to mesenchymal transition marker *twist1b*, along with fibrotic-response genes periostin-b (*postnb*) and collagen isoforms such as *col12a1b*. In contrast, the zEP cells express cell-adhesion proteins, including *cldnc* and *jam2b*, distinct from other epicardial cell types. Cells in the cEP instead maintain *tcf21/tbx18* expression without expressing these functional markers (Figure 4D / Figure S4B).

Having identified activated fibroblast-like cells from both endothelial and epicardial compartments, we next investigated how these cell populations respond to injury in both species. We first calculated the proportion of endothelial cells in the fEC clusters in each sample. Both zebrafish and medaka display a 2-3 fold increase in fEC cell proportion at 3 d.p.i., and a return to baseline levels by 14 d.p.i. (Figure 4E). We next investigated the factors that define the individual fEC clusters. Among these cells, clusters fEC2 and fEC3 were most strongly marked by expression of cell cycle progression markers (Figure S4A). Furthermore, cell cycle scoring using Seurat v4 indicated that cells in clusters fEC2 and fEC3 are nearly uniformly in S or G2M phase, respectively (Figure S4D). These proliferating fibroblast-like cells also increased in proportion after injury in both species and returned to normal levels by 14 d.p.i. (Figure 4F). Among epicardial cells, we observed a similar increase in the proportion of collagen-producing fEP cells in both species (Figure 4G). Thus, both zebrafish and medaka employ a strategy of fibroblast activation from both endothelial and epicardial compartments after injury, with a shared proliferative response in endothelium-derived cells that contribute to collagen production and scar formation.

We know relatively little about the makeup and dynamics of cardiac scar tissue between regenerating and non-regenerating species. To investigate whether zebrafish might have a unique scar composition, we examined expression of all collagen isoforms detected in over 150 cells in the dataset. It has been observed previously that zebrafish epicardial-derived fibroblasts upregulate collagen XII isoforms after injury, and that these populations promote regeneration^13, 40^. We observe a strong upregulation in expression and proportion of epicardial cells expressing collagens *col12a1a* and *col12a1b* in zebrafish, but a substantially more muted upregulation in medaka (Figure S4E). In addition, while most collagens we examined were upregulated in both species after injury, zebrafish epicardial and endothelial cells upregulated collagen V and VI isoforms not seen in medaka (Figure S4E). Matrix metalloproteinases (mmps) degrade collagen matrices and have been proposed to play important roles during regeneration and scar clearance^41, 42^. We compared expression levels of all expressed mmp genes and found shared upregulation of several mmps in endothelial cells of both species, but also species-specific upregulation of *mmp15/16* in zebrafish and *mmp19* in medaka (Figure S4F).Thus, while aspects of the fibrotic response program are shared between species, zebrafish fibroblasts upregulate a subset of collagens and matrix remodeling factors distinct from medaka. It remains to be determined what role, if any, these collagen and mmp isoforms play during regeneration.

### Medaka epicardial and endothelial cells fail to produce many pro-regenerative signals

Endothelial and epicardial cells are a major source of signals that promote revascularization and cardiomyocyte proliferation after cardiac injury in zebrafish^43, 44^. To determine which of these signals are unique to the regenerating heart, we examined expression patterns for four well-characterized pro-regenerative signals: neuregulin-1 (*nrg1*), retinoic acid synthesizing enzyme *aldh1a2*, ciliary neurotrophic factor (*cntf*), and the chemokine *cxcl12a.* Examination of their expression patterns revealed that each of the four genes were most predominantly expressed in epicardial and endothelial cells (Figure 5A).

**Figure 5:**
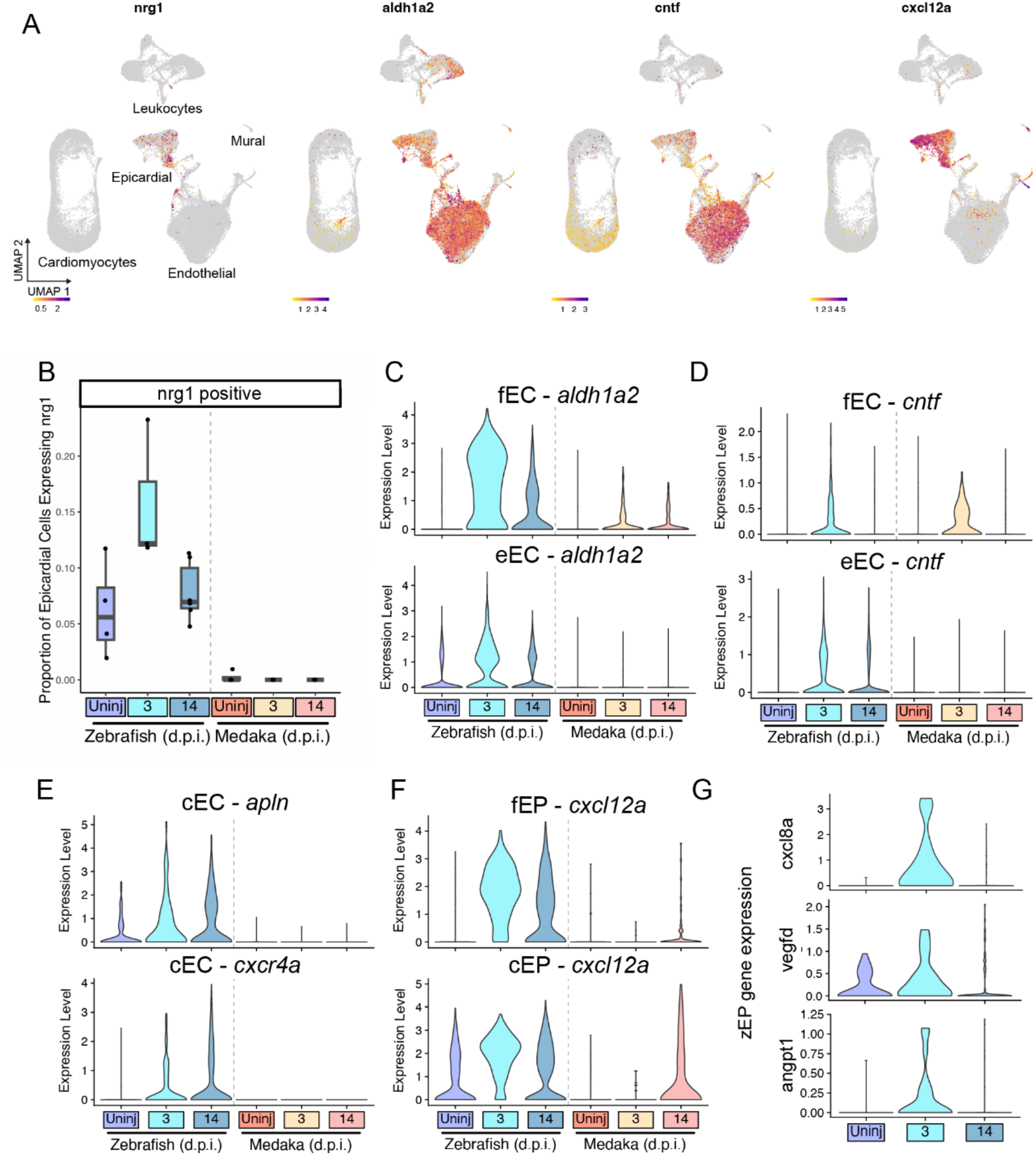
Medaka epicardial and endothelial cells fail to produce many pro-regenerative signals. (A) Gene expression feature plots for the indicated pro-regenerative genes in all cells. (B) Quantification of the proportion of all epicardial cells expressing *nrg1* in the indicated species and time point (C-G). Gene expression violin plots showing induction of pro-regenerative signals in the indicated species and time point comparing: (C) *aldh1a2* expression in fEC and fEP cells, (D) *cntf* expression in fEC and eEC cells, (E) *cxcl12a* expression in fEP and cEP cells, (F) *cxcr4a* and *apln* expression in cEC cells, and (G) *cxcl8a*, *vegfd*, and *angpt1* expression in zEP cells.

The secreted peptide nrg1 influences cellular survival and proliferation through interactions with epidermal growth factor receptors^45^. After cardiac injury, *nrg1* has been shown to be upregulated in epicardial-derived cells of the zebrafish heart and is sufficient to promote cardiomyocyte proliferation and cardiac hypertrophy, indicating a critical role in regeneration^14^. We examined *nrg1* expression patterns and found that it is mostly restricted to epicardial cells and a small number of endothelial cells (Figure S5A). We quantified the number of *nrg1+* epicardial cells and found a ∼2-fold increase at 3 d.p.i. in zebrafish, returning to base levels by 14 d.p.i. In contrast to zebrafish, we detected few *nrg1*+ medaka epicardial cells in any condition (Figure 5B). The lack of *nrg1* expression from the medaka epicardium may partially explain a lack of cardiomyocyte proliferation and regeneration.

Signaling by retinoic acid (RA), synthesized in zebrafish by the enzyme aldh1a2, is critical for development and regeneration across many types of organs and tissues^46^. In zebrafish, injury-induced RA synthesis in the epicardium and endocardium is indispensable for cardiac regeneration^15^. We observed *aldh1a2* expression in both epicardial and endothelial cells (Figure 5A / Figure S5B). Among endothelial cells, *aldh1a2* was strongly induced among the zebrafish fibroblast-like fEC at 3 d.p.i., and to a lesser extent in medaka fEC. Zebrafish eEC cells also upregulated *aldh1a2* after injury, but this response was not observed in medaka (Figure 5C), consistent with prior reports that failed to detect endocardial RA synthesis response in medaka^20^. Signaling by *cntf* has a well-established role in neural regeneration, and has recently been shown to enhance ventricular regeneration when exogenously supplied to zebrafish^47, 48^.

We find evidence for endogenous *cntf* expression predominantly in endothelial cells (Figure S5C). Similar to *aldh1a2* expression patterns, we observe a strong induction of *cntf* in fEC cells in both zebrafish and medaka, but expression of *cntf* among eEC cells was only observed in zebrafish (Figure 5D). Together, these results suggest that the endocardium of medaka, the most abundant cell type in the heart, fails to mount a systemic pro-regenerative response to injury as seen in zebrafish.

The zebrafish ventricle rapidly revascularizes after injury, but medaka have limited re-growth of blood vessels after injury^20^. This process is critical for regeneration, and evidence suggests that nascent blood vessels provide a scaffold for regrowing cardiomyocytes during regeneration^33^. We investigated whether there was evidence of revascularization through analysis of *apelin* and *cxcr4a* expression, which are both upregulated in growing coronary vessels^33^. Consistent with our understanding of previous literature, both *apln* and *cxcr4a* were elevated in zebrafish cEC cells after injury, but remained low at all time points in medaka (Figure 5E). Cxcl12a peptide acts as a ligand for cxcr4a, promoting angiogenesis and regeneration^47^. We observed that expression of *cxcl12a* was mostly restricted to the epicardial compartment (Figure 5A/S5D). Following injury, *cxcl12a* was highly upregulated in fEP and cEP cells in zebrafish by 3 d.p.i., but only apparent by 14 d.p.i. In medaka (Figure 5F). These results indicate that the zebrafish epicardium rapidly releases pro-angiogenic signals that promote rapid revascularization unique to this regenerating species.

When examining epicardial cells, we identified a small number of zebrafish-specific cells in the zEP cluster. In contrast to the fEP, the zEP cluster is only composed of zebrafish cells and does not change in abundance after injury (Figure S5E). These cells express epicardial marker *tcf21* and appear to have a unique injury-response profile. At 3 d.p.i., we observe an increased level of expression of pro-angiogenic signaling in zEP, including factors *angpt1, vegfd*, and *cxcl8a*, each known to play a role in promoting angiogenesis^50–52^ (Figure 5G). Thus, while more evidence is needed to determine whether the zEP cells are truly unique to zebrafish, they represent a potential hub of pro-angiogenic signaling that may promote revascularization after injury in zebrafish.

### Medaka lack primordial myocardium and have few compact layer cardiomyocytes

Although medaka and zebrafish hearts are similar in overall morphology, they may not share similar populations of cardiomyocytes. It has been proposed that some cardiomyocyte populations might have outsized contributions to the pool of regenerating cells^7^. The zebrafish myocardium contains three main types of cardiomyocytes with distinct spatial arrangements: the inner mass of trabecular cardiomyocytes (tCMs), the outer layer of cortical cardiomyocytes (cCMs), and a single-cell-thick layer of primordial cardiomyocytes that sits underneath the cortical layer^53^. When comparing ventricle sections, we noticed a distinctly thin cortical layer in the medaka ventricle compared to zebrafish (Figure 6A). The cortical myocardium has been proposed to play an important role in regeneration^54^, yet some types of fish do not have this layer^55^. We further compared the composition of the myocardium by reclustering cells expressing cardiomyocyte markers *myl7* and *actc1a*, identifying 4 cell clusters (Figure 6B). We identified one cluster of tCMs, which express canonical CM markers but lack expression of transcription factor *tbx5a*^56^. The remaining cell clusters expressed transcription factors *tbx5a*, as well as *gata6* and *mef2d*, and are assigned as cCMs (Figure 6C). Comparing the cellular compositions of each cardiomyocyte type revealed a skewing toward trabecular cardiomyocytes in medaka, and fewer cells clustering with cortical CMs, consistent with our imaging observations (Figure 6D).

**Figure 6:**
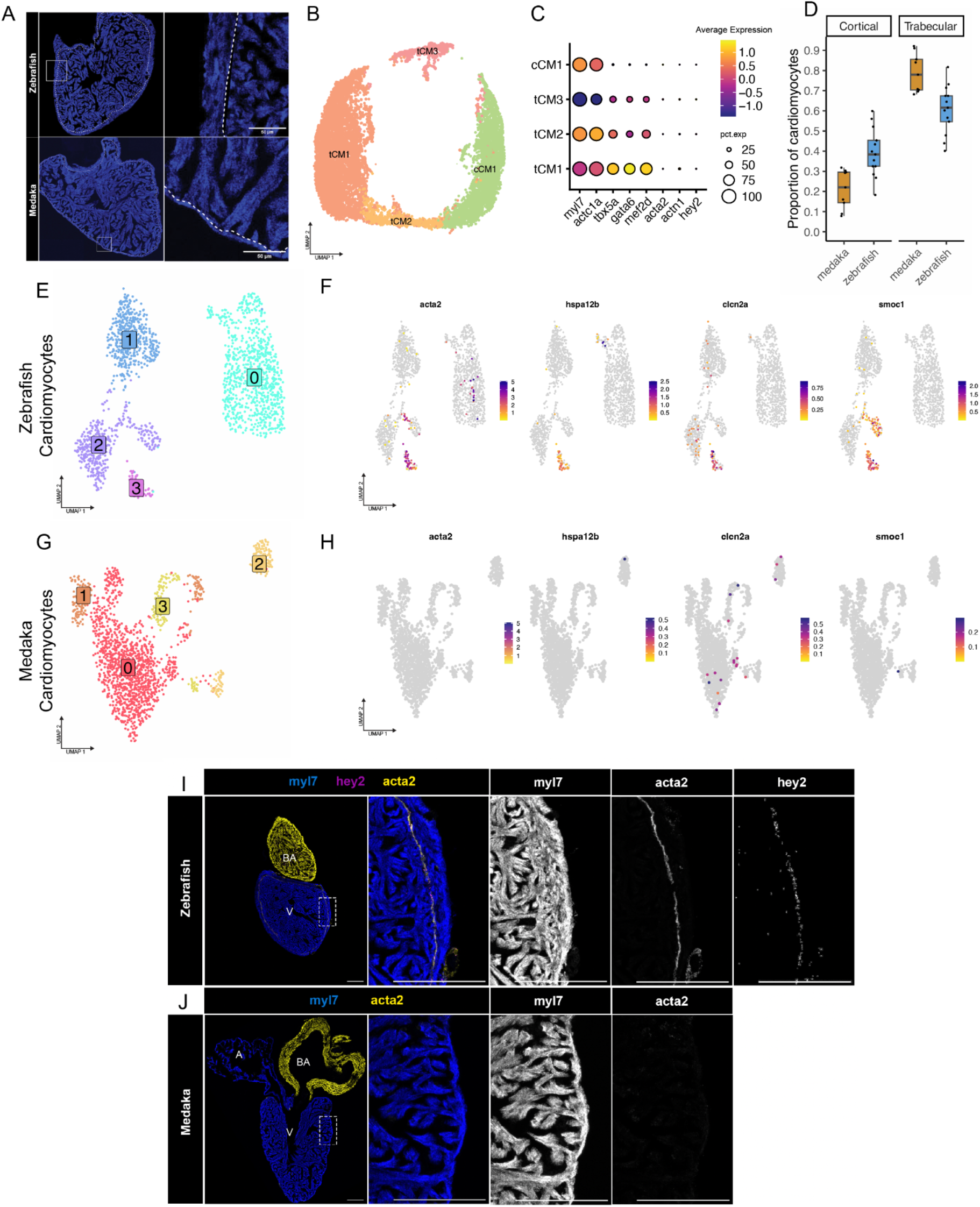
Medaka lack primordial myocardium and have few compact layer cardiomyocytes. (A) RNA in situ hybridization of myl7 labeling myocardium in zebrafish and medaka ventricles. Dotted line indicates border between trabecular and cortical layers. (B) UMAP embedding of re-clustered cardiomyocytes identified as either trabecular (tCM) or cortical (cCM). (C) Gene expression dot plot showing expression of marker genes for each CM cell cluster. (D) Proportion of cardiomyocytes in trabecular or cortical cell clusters from all single-cell samples from zebrafish and medaka. (E-F) UMAP embedding of ventricular cardiomyocytes clustered separately from uninjured (E) zebrafish or (F) medaka. (G-H) Gene expression feature plots for top marker genes for primordial cardiomyocytes in zebrafish (E) or medaka (F). (I) RNA in situ hybridization of *myl7*, *acta2*, and *hey2* in uninjured zebrafish hearts (scale = 200µM). (J) RNA in situ hybridization of *myl7* and *acta2* in uninjured medaka heart (scale = 200µM)

Primordial cardiomyocytes are of interest as a potential source for newly regenerated myocardium due to their undifferentiated morphology and embryonic-like gene signature^57, 58^. Initially, we did not observe a cell cluster expressing markers for the primordial myocardium (*acta2/hey2/actn1*) in the integrated single-cell object (Figure 6B). However, species-specific clustering of uninjured cardiomyocytes revealed a small cluster of zebrafish cells expressing primordial CM makers, including *acta2*, *hspa12b, clcn2a,* and *smoc1* (Figure 6E-F). In contrast, clustering of uninjured medaka CMs failed to identify any group of cells expressing analogous cell markers (Figure 6G-H). We examined the spatial patterns of top primordial CM markers *acta2* and *hey2* expression in the zebrafish ventricle and found co-localization in a thin sub-cortical cell layer of cardiomyocytes expressing both genes (Figure 6I). We searched for a similar arrangement of CMs in medaka ventricles by probing for *acta2* expression (*hey2* is not annotated in the medaka genome). Imaging revealed no evidence of either *acta2+* cardiomyocytes or an analogous primordial layer in medaka (Figure 6J). Together, our transcriptomic and imaging data suggest that medaka lack primordial cardiomyocytes and have a diminished or absent compact myocardium, adding another factor that may influence the ability to regenerate.

### Zebrafish cardiomyocytes share a cardioprotective gene signature with neonatal mouse

Neonatal mammals are capable of heart regeneration but retain permanent scars if injured as adults. To investigate whether the neonatal mammal heart might share similarities with zebrafish, we compared injury responses in our dataset to a recently published single-nucleus RNA-seq dataset that assessed cardiomyocyte responses to injury in regenerating postnatal day 4 (p4) mice and non-regenerating postnatal day 11 (p11) mice^59^. We used differential gene expression testing to identify all upregulated genes at 3 d.p.i. in cardiomyocyte cells in the p4 and p11 mice as well as in zebrafish and medaka cardiomyocytes. Using orthology assignments from Ensembl, we determined which upregulated genes with one-to-one orthology were shared or unique in each species. Indeed, zebrafish cardiomyocytes shared three times as many injury-upregulated genes with P4 mouse (33 genes) compared to the non-regenerating P11 mouse (11 genes) (Figure 7A). In contrast, Medaka cardiomyocytes shared more genes in common with the p11 mouse (12 genes) (Figure 7B). Remarkably, a large proportion of the shared injury-response genes between zebrafish and p4 mouse cardiomyocytes have well-described cardioprotective roles, promoting cardiomyocyte survival and guarding against metabolic stress (Table S1). Examination of gene expression of these cardioprotective genes revealed a conspicuous lack of induction in both medaka and p11 cardiomyocytes (Figure 7C). These results indicate that regenerating vertebrates deploy an evolutionarily-conserved cardioprotective program immediately after myocardial injury.

**Figure 7:**
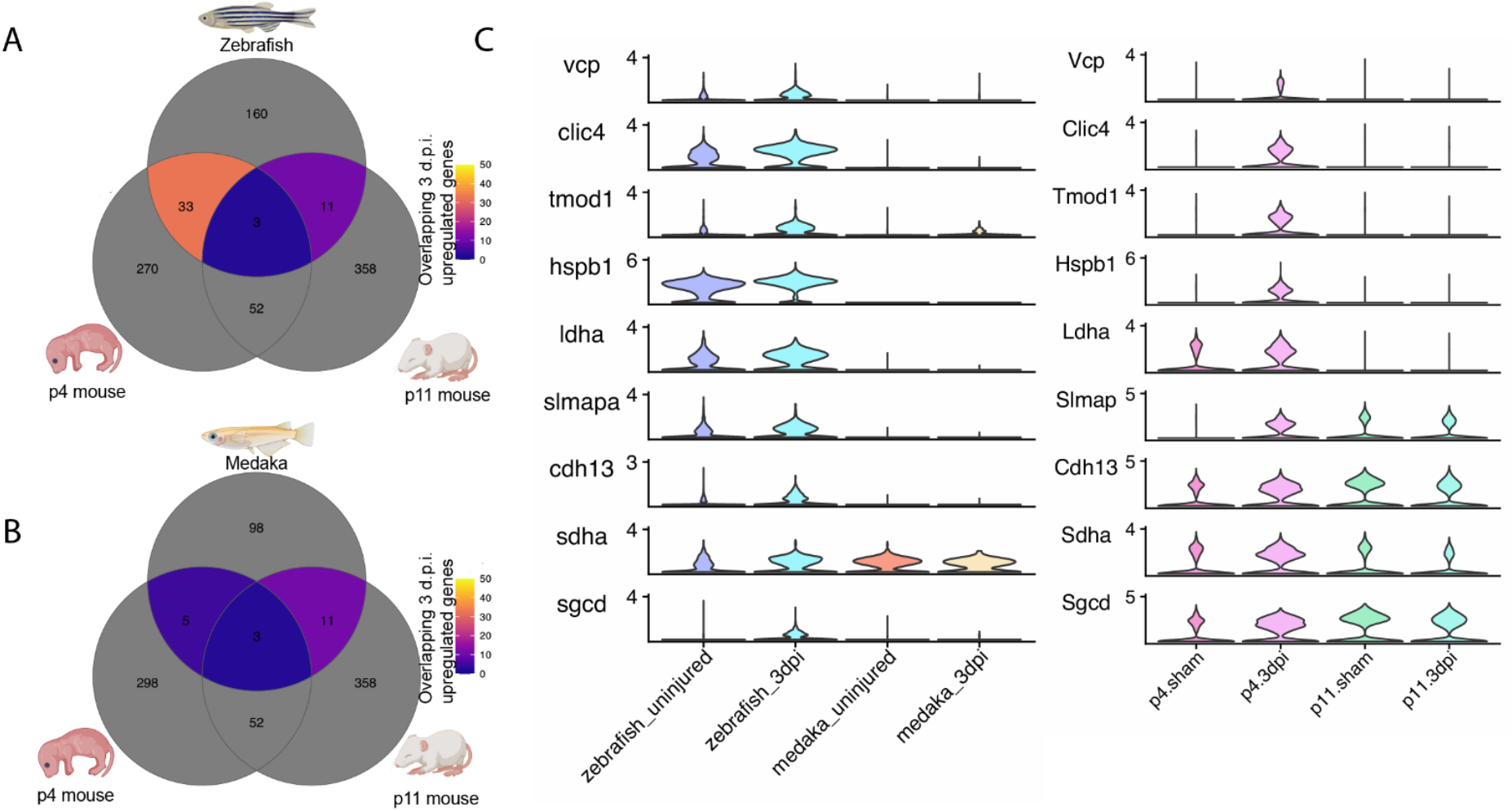
Zebrafish cardiomyocytes share a cardioprotective gene signature with neonatal mouse. (A-B) Venn diagram counting overlapping genes upregulated at 3 d.p.i. in postnatal day 4 or 11 mouse cardiomyocytes with (A) zebrafish cardiomyocytes or (B) medaka cardiomyocytes. (C) Gene expression violin plots of cardioprotective genes uniquely upregulated in regenerating p4 mouse and zebrafish cardiomyocytes.

## DISCUSSION

In this study, we used single-cell RNA sequencing to compare the injury responses of zebrafish and medaka heart ventricles. Our analysis revealed several previously undescribed differences between the two species, including variations in immune cell populations, endothelial and epicardial cell signaling, and the composition of the myocardium itself. In addition, our results corroborated and expanded upon previous observations in medaka, including altered macrophage recruitment and dampened retinoic acid signaling^20, 22^. These findings highlight the power of comparative scRNAseq to identify species-specific differences in biological processes and have generated hypotheses for future investigations into the mechanisms of cardiac regeneration in these species.

Our comparative analysis of immune cell populations and gene expression in response to injury revealed major differences between zebrafish and medaka. Previous comparisons showed that the hearts of these two species express a different repertoire of immune genes after injury^22^. Remarkably, injection of poly(I:C), a potent activator of interferon signaling, can promote cardiac revascularization and regeneration in medaka. Interferon signaling seems to have pleiotropic effects depending on the tissue type, where it is deleterious in some cases but promotes regeneration in others^31, 32^. Our findings indicate a major difference between zebrafish and medaka in response to injury is the presence of endogenous interferon signaling specific to zebrafish. Interferon signals may play a role in modulating immune cell behaviors, perhaps explaining why zebrafish appear to recruit macrophages to the ventricle injury site while medaka do not. Additionally, while pro-regenerative genes such as *aldh1a2* or *nrg1* are not canonical ISGs, it remains to be determined whether indirect effects of interferon might promote their expression. Indeed, we find that the endocardium and epicardium, which are the main source of pro-regenerative signaling, exhibit the most robust ISG expression. Experimentally studying the role of endogenous interferon signaling during heart regeneration represents a compelling direction for future inquiry.

Our examination of endothelial and epicardial cell populations uncovered several injury-response behaviors that are shared between zebrafish and medaka. Overall, both species shared a similar pattern of fibroblast activation from both the endothelial and epicardial compartments. Our comparison of the set of collagen and matrix remodeling factors made by each species showed a mostly overlapping pattern of induction after injury. This included collagen XII isoforms in epicardial-derived fibroblasts, which were upregulated in both species after injury, but to a higher degree in zebrafish. These collagen XII-producing fibroblasts are of special interest as they are required for proper regeneration in zebrafish^13^. We also found evidence of some species-specific matrix-remodeling proteins and collagen isoforms, warranting further investigation into whether the makeup of the scar itself may promote regeneration in zebrafish.

While we observed similar strategies of fibroblast activation from the endothelial and epicardial compartments in both species, medaka failed to produce critical signals required for regeneration from these cells. In particular, medaka fibroblasts expressing epicardial markers had a conspicuous lack of *nrg1* expression, which itself is sufficient to activate regenerative programs in the heart^14, 69^. Interestingly, we observed a post-injury upregulation of some pro-regenerative signals in medaka endothelial-like fibroblasts, including *aldh1a2* and *cntf*. These same signals were highly upregulated in zebrafish endocardial clusters, but notably absent from medaka endocardium. In zebrafish, the endocardium undergoes systemic morphological changes in response to injury, coincident with strong upregulation of *aldh1a2* and other signaling factors even in cells distal from the wound area^15^. Further experiments are needed to define the specific stimuli that trigger this systemic response and to determine why medaka endocardial cells appear to be less responsive to injury.

Finally, we investigated the cellular makeup of the myocardium in zebrafish and medaka. Our results reveal marked differences in the cardiomyocyte populations of these two species, which are discernible even in non-injured samples. Specifically, we found that medaka have a highly diminished or absent compact myocardium, as evidenced by our single-cell sequencing and imaging analyses. This observation is notable, given that previous studies have indicated that the compact myocardium is activated after injury in zebrafish and may serve as a source of pro-regenerative cardiomyocytes^54^. Additionally, we were unable to identify primordial layer cardiomyocytes in medaka using either single-cell gene expression analysis or by imaging of key markers. However, we did observe that the primordial myocardium in zebrafish expresses *acta2*, in conjunction with *hey2*, an embryonic cardiomyocyte marker, indicating an immature transcriptional state^70^ . These findings are especially interesting in light of recent studies demonstrating that *acta2+* embryonic-like cardiomyocytes are present in neonatal mammalian hearts but are lost as the animal ages, coincident with the decline of regenerative capacity^59, 71^. Thus, the presence of immature cardiomyocytes in uninjured hearts is notably correlated with regenerative capability. Further investigation is warranted to explore whether primordial layer cardiomyocytes are required for regeneration.

Medaka and zebrafish have been used extensively as laboratory model vertebrates, and have similar care requirements and body plans. While it was known that medaka are incapable of heart regeneration, it could have been assumed that medaka as fellow teleost fish would only have minor differences in cardiac structure. Our study found a surprising number of distinguishing characteristics of the medaka heart compared to zebrafish. These differences included not only changes in cellular behaviors but also substantial changes in the structure of the myocardium itself. Given the number of notable differences we observed between zebrafish and medaka, a wider phylogenetic survey of cardiac injury responses will be particularly useful to identify features that correlate with heart regeneration or non-regeneration across the teleost phylogeny. These observations highlight how biodiversity within shorter evolutionary distances can enable comparative studies that reveal fundamental insights about the gain or loss of complex traits.

## METHODS

### Fish husbandry

Wild-type Tübingen zebrafish and CAB medaka, aged 6-18 months, were used for all experiments. All zebrafish and medaka work was performed at University of Utah’s CBRZ zebrafish facility. This study was conducted under the approval of the Office of Institutional Animal Care and Use Committee (IACUC no. 20-07013) of the University of Utah’s animal care and use program.

### Cardiac Cryoinjury

Cryoinjuries were performed on the ventricle apex of both medaka and zebrafish as described previously^8^. Briefly, 0.02% and 0.04% Tricaine (MS-222) was used to anesthetize zebrafish and medaka, respectively. Fish were mounted on a moist sponge. The ventricle apex was exposed by making a small thoracic incision using forceps and dissecting scissors. A cryoprobe was constructed as described previously using a 0.5 mm diameter copper wire^72^. After submersion in liquid nitrogen for at least 2 minutes, the probe was placed in contact with the ventricle apex for exactly 23 seconds. After injury, fish were placed back into freshwater tanks to recover, then transferred back into the fish facility for monitoring.

### Histology

Hearts were fixed in 4% paraformaldehyde overnight, then sucrose treated and embedded in Optimal Cutting Temperature (O.C.T.) embedding medium. 6 µm coronal sections of the heart were cut using a Leica Cryostat. Bouin’s solution was used to fix heart tissue before staining with PT/PM (Phosphotungstic Acid/Phosphomolybdic Acid) and AFOG (Methyl Blue, Orange G, and Acid Fuchsin) stain. After staining, heart tissue was washed with 0.5% acetic acid, 100% ethanol, and xylenes. Fibrinogen and collagen deposition from heart sections with representative injuries for both medaka and zebrafish were chosen to demonstrate comparative injury phenotypes at 3 and 14 days post-injury. Measurements were made using Fiji, ImageJ, software.

### Single-cell isolation and library preparation

To prepare cells for scRNA-seq, adult fish were euthanized by immersion in ice-cold water. After dissection and removal of hearts, the remaining atrium and bulbus arteriosus tissue were cut away with scissors to isolate the ventricle. Ventricles were placed in PBS solution and gently squeezed with forceps to remove residual blood prior to digestion. Whole ventricles were placed in 200 µl dissociation solution containing 1mg/mL liberase DH (Millipore Sigma cat# 5401089001) in 1X HBSS. Ventricles were digested at 37°C for 30 min on a benchtop shaker at 250RPM with additional pipetting every 5 min to break apart the tissue. After complete digestion, the reaction was quenched with ice-cold PBS/50% Fetal Bovine Serum and passed through a 40 µM strainer. The cells were then centrifuged at 250 RCF for 10 minutes at 4°C and resuspended in ice-cold PBS with 0.04% Bovine Serum Albumin. Cells were then assessed for viability and dissociation quality using acridine orange and propidium iodide staining with a Nexcelom Cellometer automated cell imager. Each sample viability was assessed to be >85%. Cells were loaded onto a 10X Genomics Chromium controller and processed according to the manufacturer’s specifications. cDNA libraries were prepared for sequencing on the Illumina Novaseq platform at a depth of approximately 400M reads/sample.

### Data integration and clustering

Single-cell RNA seq reads were processed with Cellranger from 10X Genomics with the default settings, using GRz11 v4.3.2 reference for zebrafish read alignments and ASM223467v1 for Medaka. Gene expression matrices were imported and processed in Seurat v4 for subsequent processing. A preliminary filtering step was used to remove cells with less than 200 unique RNA features or cells with more than 40% mitochondrial RNA. To cluster cells across species and correct for batch effects, we used the data integration function in Seurat v4 which uses gene anchors with shared names to apply correction vectors to correct for batch and species effects and can allow for cross-species integration^23^. To maximize the number of usable anchors we renamed 596 medaka genes that were confidently assigned as one-to-one orthologs as determined by Ensembl orthology assessment. An initial round of clustering was performed on log normalized gene expression, and erythroid cells (*hbba1/hbaa1* positive) and platelets *(thbs1b/itga2b* positive) were filtered out. An additional filtering criterion was then applied to remove non-myocyte cells with greater than 15% mitochondrial RNA. The filtered cells were then re-clustered and annotated by marker gene expression.

### RNAscope in situ hybridization

Prior to performing RNAscope (Advanced Cell Diagnostics, Hayward, CA), fish hearts were fixed overnight in 4% PFA in 1x PBS, sucrose treated, embedded in OCT, and cryosectioned into 12 µm coronal sections. RNAscope® Multiplex Fluorescent Detection Kit v2 protocol was completed according to the manufacturer’s protocol for standard fixed-frozen tissue samples. See key resources table for probe details.

### Imaging

Brightfield images of AFOG stains were acquired with the Zeiss Axio Scan Z1 instrument. Fluorescence images were captured on a Zeiss 880 Airy Scan confocal microscope. Tiling and stitching were completed by ZEN Black software. Images displayed are maximum intensity projections. Adjustments of contrast and brightness were made using ImageJ. At each step of image collection, similar settings for both species were used. When images were used for quantification, all images were collected using the same laser settings (gain, laser power, ext.).

### Imaging quantification

Images of RNAscope sections that marked cardiomyocytes (*myl7*), macrophages (*mpeg1.1*), and nuclei (DAPI) were prepared and processed with ImageJ. To distinguish between the intact ventricle and wound area, ImageJ thresholding of the *myl7* channel was completed to make two regions of interest (ROI)—a region that marked just the intact ventricle and a region that marked the intact ventricle and the wounded area. These ImageJ ROI were used to calculate the areas of each region. Then the ROIs were applied to the *mpeg1.1* and DAPI channels to mask either the wound area or the intact ventricle. Masked images were uploaded to CellProfiler software (version 4.2.1) using a modified CellProfiler “Speckle Counting” pipeline^72^. Nucleated cells were identified using the DAPI channel and the IdentifyPrimaryObjects CellProfiler module. *mpeg1.1* speckles were identified using the *mpeg1.1* channel and the IdentifyPrimaryObjects CellProfiler module. Objects were related using the RelateObjects CellProfiler module. Cells were identified as macrophages when a *mpeg1.1* speckle fell within a mask created by the DAPI channel.

### Key resources table

| Reagent type or resource | Designation | Source or reference | Identifiers | Additional information |
| --- | --- | --- | --- | --- |
| Commercial assay, kit | RNAscope® Multiplex Fluorescent Detection Kit v2 | Advanced Cell Diagnostics | Ref: 323110 |  |
| Sequence-based reagent | RNAscope Probe - Dr-myl7 | Advanced Cell Diagnostics | Ref: 455661 |  |
| Sequence-based reagent | RNAscope Probe - Ol-myl7-C1 | Advanced Cell Diagnostics | Ref: 1135361-C1 |  |
| Sequence-based reagent | RNAscope Probe - Dr-hey2-C2 | Advanced Cell Diagnostics | Ref: 508571-C2 |  |
| Sequence-based reagent | RNAscope Probe - Dr-kdrl-C2 | Advanced Cell Diagnostics | Ref: 416611-C2 |  |
| Sequence-based reagent | RNAscope Probe - Ol-LOC100049480-C2 | Advanced Cell Diagnostics | Ref: 1225041-C2 |  |
| Sequence-based reagent | RNAscope Probe - Dr-isg15-C3 | Advanced Cell Diagnostics | Ref: 559261-C3 |  |
| Sequence-based reagent | RNAscope Probe - Ol-LOC101167545-C3 | Advanced Cell Diagnostics | Ref: 1225061-C3 |  |
| Sequence-based reagent | RNAscope Probe - Dr-mpeg1.1-C2 | Advanced Cell Diagnostics | Ref: 536171-C2 |  |
| Sequence-based reagent | RNAscope Probe - Dr-LOC101169041-C2 | Advanced Cell Diagnostics | Ref: 1225071-C2 |  |
| Sequence-based reagent | RNAscope Probe - Dr-lck-C2 | Advanced Cell Diagnostics | Ref: 529951-C2 |  |
| Sequence-based reagent | RNAscope Probe - Ol-lck-C2 | Advanced Cell Diagnostics | Ref: 1135371-C1 |  |
| Sequence-based reagent | RNAscope Probe - Dr-acta2-C3 | Advanced Cell Diagnostics | Ref: 508581-C3 |  |
| Sequence-based reagent | RNAscope Probe - Ol-acta2-C3 | Advanced Cell Diagnostics | Ref: 1153791-C3 |  |
| Other | Opal 520 Reagent | Akoya Biosciences | Product #: OP-001001 | 1:1500 |
| Other | Opal 570 Reagent | Akoya Biosciences | Product #: OP-001003 | 1:1500 |
| Other | Opal 690 Reagent | Akoya Biosciences | Product #: OP-001006 | 1:750 |
| Other | Superfrost Plus Microscope Slides | Fisher Scientific | Catalog #: 12-550-15 |  |
| Peptide, recombinant protein | Liberase™ DH (Dispase High) Research Grade | Roche | Product #: LIBDH-RO |  |

## AUTHOR CONTRIBUTIONS

C.M.C. and J.A.G. conceived of the project. C.M.C. performed scRNA-seq and data analysis and wrote the manuscript. C.M.C., H.L.H., and A.V.S. performed cryosurgery procedures and animal husbandry. H.L.H. carried out in situ hybridization and imaging quantification. H.L.H. and A.V.S completed imaging. A.V.S. performed histology experiments. J.A.G. supervised the project. All authors reviewed and edited the manuscript.

## ACKNOWLEDGEMENTS

We acknowledge Cell Imaging Core at the University of Utah for the use of equipment (Zeiss 880 Airy Scan, Axio Scan.Z1, and Leica Cryostat) and thank Xiang Wang for assisting our imaging work. We thank the University of Utah high throughput genomics core and Opal Allen for assistance with single-cell sequencing. C.M.C. is supported by the National Institutes of Health grant number F32HL156644. H.L.H. is supported by the University of Utah (U of U) Undergraduate Research Opportunities Program, U of U ACCESS Scholars Program, U of U Biology Research Scholars Program, Francis Family Foundation Undergraduate Research Scholarship, and the U of U Summer Program for Undergraduate Research. Work in the Gagnon laboratory was supported by National Institutes of Health grant R35GM142950, a pilot grant from the NHLBI CvDC Bench to Bassinet program, and startup funds from the University of Utah. We thank Mike Shapiro and Chelsea Herdman for their helpful comments and review of the manuscript.

## DATA AVAILABILITY

Raw sequencing data are being submitted to GEO. A web application for user-friendly data exploration and analysis of the single-cell datasets, alongside gene expression matrices, processed Seurat objects, and R code for data processing and data visualization can be accessed at https://github.com/clay-carey/medaka_zebrafish_regeneration.

