## Supplemental Figures and Tables for "Distinct features of the regenerating heart uncovered through comparative single-cell profiling"

**Figure S1: related to Figure 1**

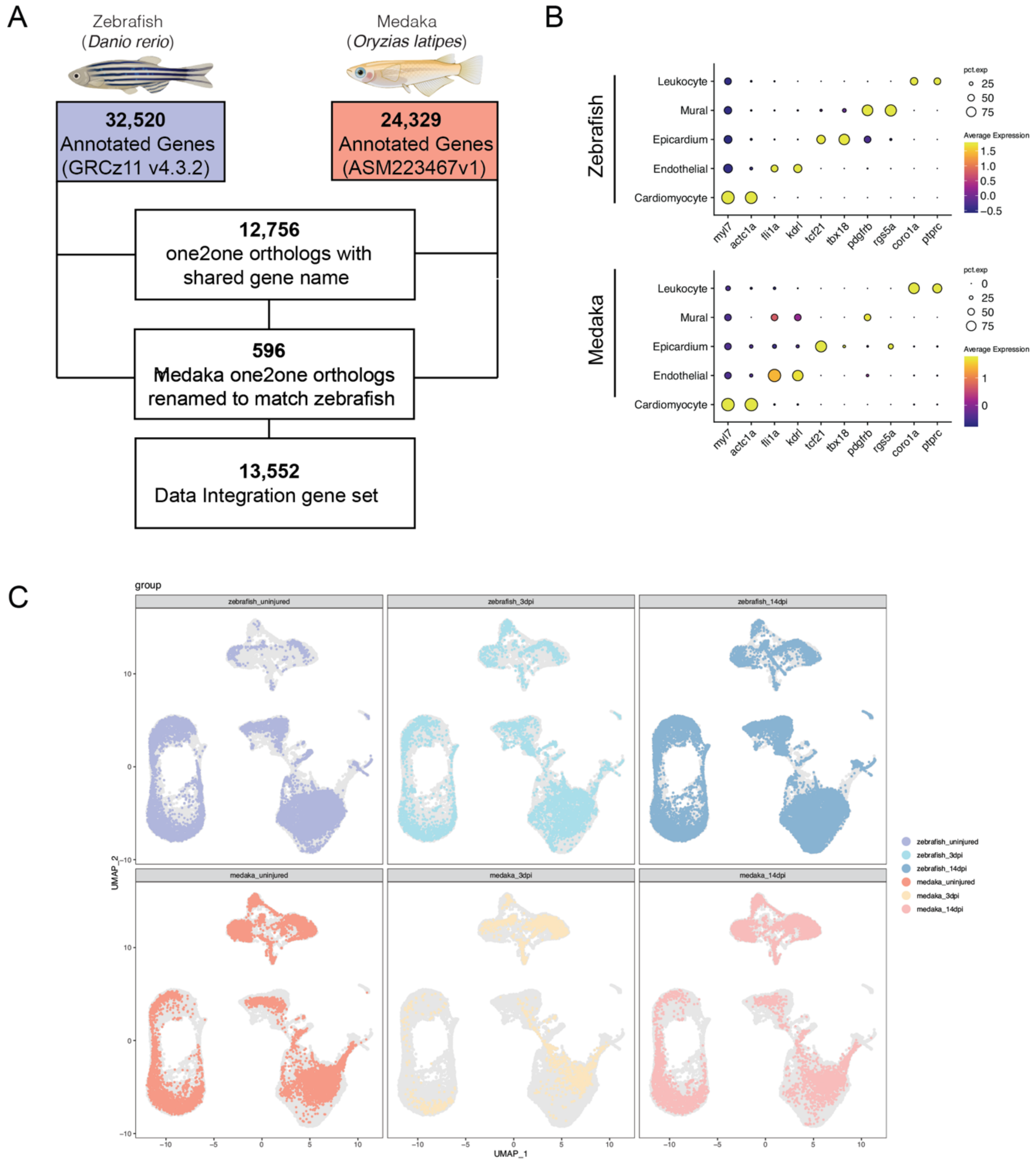


**Figure S1: A single-cell atlas of cardiac injury response in zebrafish and medaka.** (A) Overview of data integration scheme. (B) Dot plot of cardiac cell type marker gene expression split by species. (C) UMAP embeddings showing distribution of each group of samples.

### Figure S2: Related to Figure 2

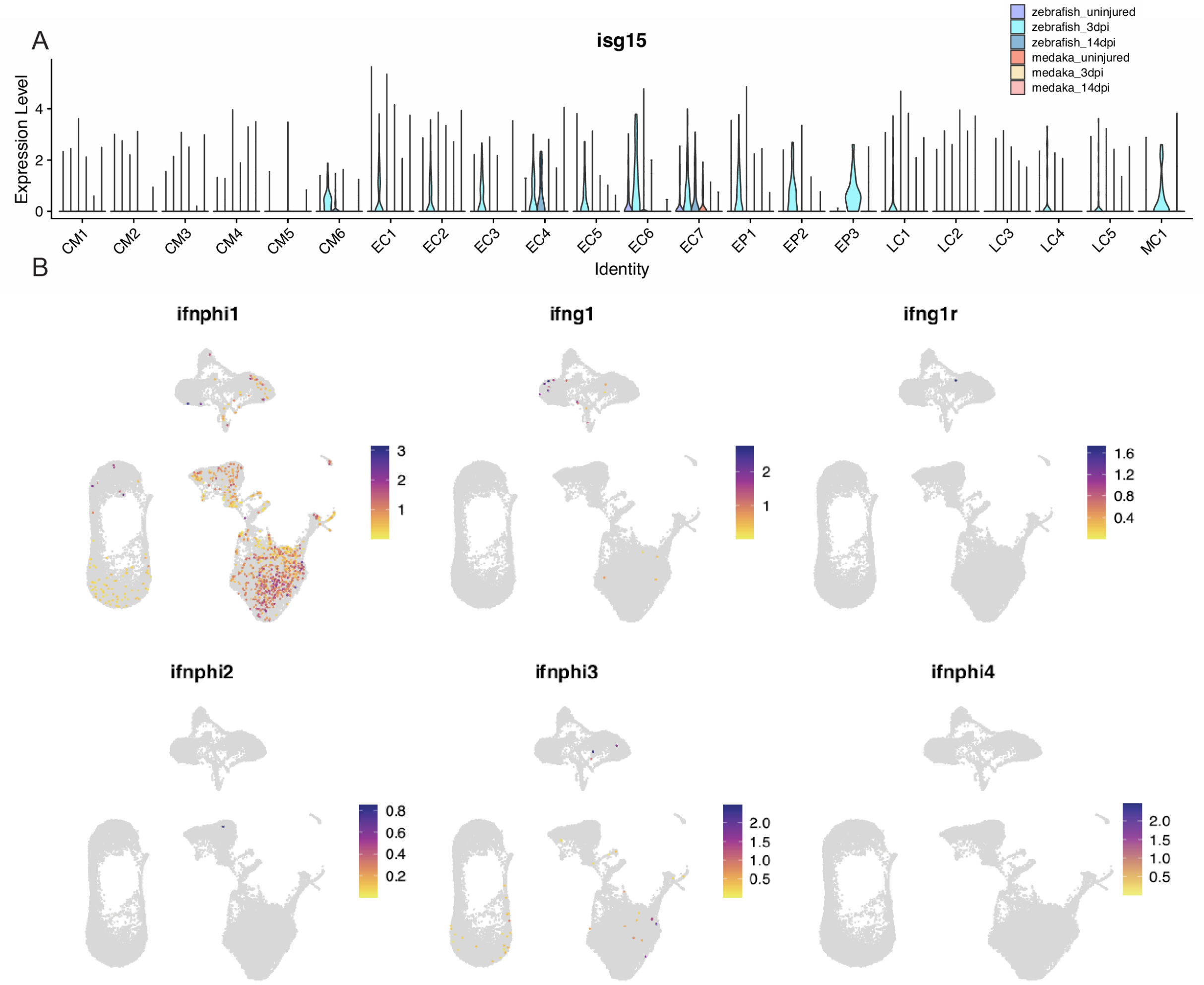


**Figure S2: Medaka lack an endogenous injury-induced interferon response.** (A) Gene expression violin plot for *isg15* in each cell cluster, grouped by species and injury status. CM = cardiomyocyte, EC = endothelial cell, EP = epicardial cell, LC = leukocyte, MC = Mural Cell. (B) Gene expression feature plots of zebrafish cells for each annotated interferon isoform in the zebrafish genome color scale = expression level.

### Figure S4: Related to Figure 4

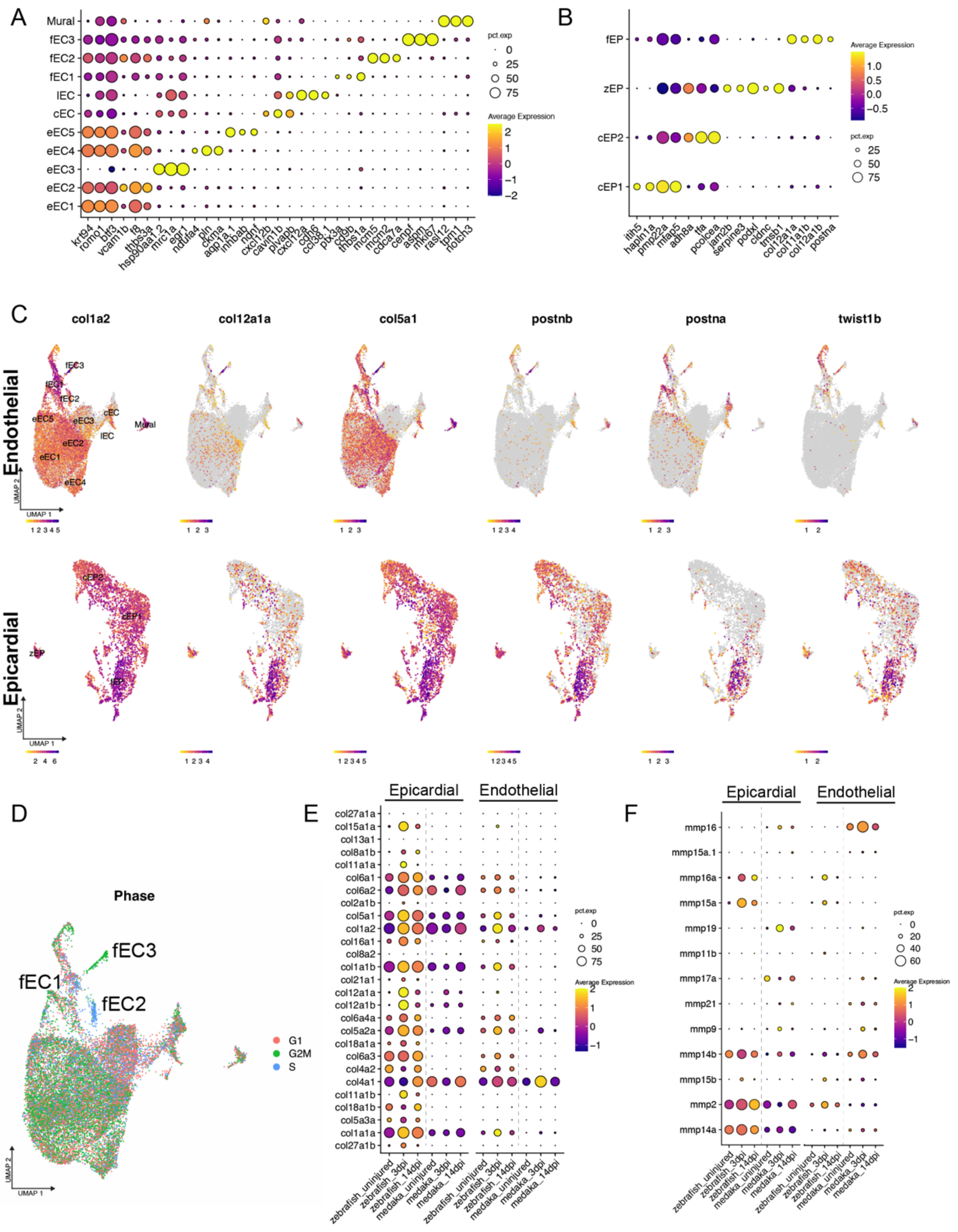


**Figure S4: Zebrafish and medaka share a partially overlapping fibrotic response to injury**
(A-B) Top markers for each individual (A) endothelial/mural or (B) epithelial cell cluster. (C) Gene expression feature plots for the indicated fibrosis genes in both epicardial (top) or endothelial (bottom) cell types.(D) UMAP embedding of endothelial cells classified by Seurat cell cycle scoring assignments highlighting uniform G2M/S phase assignments in fEC3 and fEC2, respectively. (E-F) Gene expression dot plots for all endothelial and epicardial cells for all (E) collagen and (F) matrix metalloproteinases expressed in >150 cells at the indicated time point and species.

**Figure S5: Related to Figure 5**

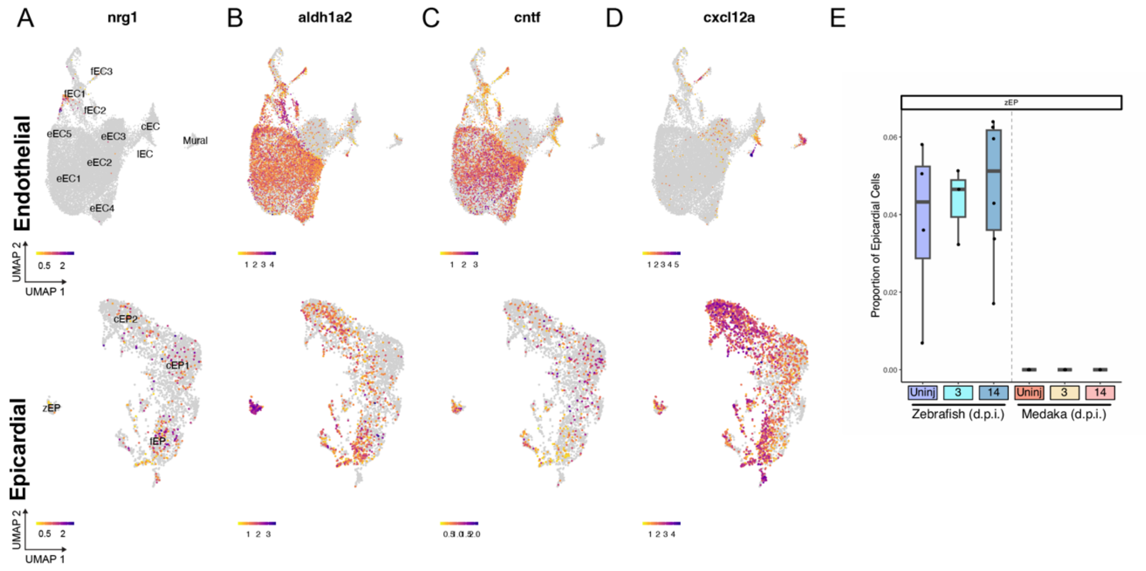


**Figure S5. Medaka epicardial and endothelial cells fail to express many pro-regenerative signals.**(A-D) Gene expression feature plots of endothelial cells (top) and epicardial cells (bottom) showing normalized gene expression of (A) *nrg1,* (B) *aldh1a2,* ( C) *cntf,* and (D) *cxcl12a* in integrated objects of both species*.* (E) Quantification of proportion of epicardial cells clustered as zEP at each time point in each species.

**Table S1: Cardioprotective gene functions, related to Figure 7**

| Gene | Effect | Citation |
|---|---|---|
| Valosin Containing Protein (*vcp*) | Promotes cardiomyocyte survival through preservation of mitochondrial integrity | Lizano *et al.* 2013[59] |
| Chloride intracellular channel protein (*clic4*) | Maintains $Ca^{2+}$ homeostasis after injury and promotes cell survival | Ponnalagu *et al.* 2022[60] |
| Tropomodulin-1 (*tmod1*) | Involved in regulating cytoskeletal dynamics after injury, promoting cardiomyocyte migration. | Poch *et al.* 2022[61] |
| Heat shock protein b1 (*hspb1*) | Protects myofibrils during cardiac stress | Brundel *et al.* 2008[62] |
| Lactate dehydrogenase A (*ldha*) | Involved in metabolic reprogramming and promotes CM proliferation | Chen et al. 2022[63] |
| Sarcolemma associated protein a (*slmapa*) | Provides mechanical support to cardiomyocyte membranes | Nader 2019[64] |
| T-cadherin (*cdh13*) | Protects against cardiac injury via interaction with adiponectin and AMPK signaling | Denzel et al. 2010[65] |
| succinate dehydrogenase complex, subunit A (*sdha*) | Upregulated in regenerating border zone regenerating cardiomyocytes, involved in metabolic reprogramming. | Spelat et al. 2022[66] |
| Sarcoglycan delta (*sgcd*) | Stabilizes cardiomyocyte membrane in response to stress via ECM interaction. | Morikawa et al. 2015[67] |
